# Paradox resolved: stop signal race model with negative dependence

**DOI:** 10.1101/158485

**Authors:** Hans Colonius, Adele Diederich

## Abstract

The ability to inhibit our responses voluntarily is an important case of cognitive control. The stop-signal paradigm is a popular tool to study response inhibition. Participants perform a response time task (*go task*) and, occasionally, the go stimulus is followed by a stop signal after a variable delay, indicating subjects to withhold their response (*stop task*). The main interest of modeling is in estimating the unobservable stop-signal processing time, that is, the covert latency of the stopping process as a characterization of the response inhibition mechanism. In the *independent race model* the stop-signal task is represented as a race between stochastically independent go and stop processes. Without making any specific distributional assumptions about the processing times, the model allows to estimate the mean time to cancel a response. However, neurophysiological studies on countermanding saccadic eye movements have shown that neural correlates of go and stop processes consist of networks of mutually *interacting* gaze-shifting and gaze-holding neurons. This poses a major challenge in formulating linking propositions between the behavioral and neural findings. Here we propose a *dependent race model* that postulates perfect negative stochastic dependence between go and stop activations. The model is consistent with the concept of interacting processes while retaining the simplicity and elegance of the distribution-free independent race model. For mean data, the dependent model’s predictions remain identical to those of the independent model. The resolution of this apparent paradox advances the understanding of mechanisms of response inhibition and paves the way for modeling more complex situations.

---

**A** recurrent theme in cognitive modeling is the difficulty to uniquely identify the processes underlying the generation of response times and probabilities in behavioral paradigms like simple yes-no tasks or same-different judgments of stimulus pairs. A prime example is the serial vs. parallel processing issue (1–3) showing that even rigorous mathematical analyses of the underlying assumptions may not always resolve the difficulty completely. Recently, efforts in model-based cognitive neuroscience to constrain behavioral models by findings obtained from neuroscientific methods have intensified, from spike train analyses to EEG and fMRI recordings (4). One area where important advances have emerged from this joint endeavor is the modeling of cognitive control and, in particular, response inhibition (5, 6). Response inhibition refers to the ability to suppress responses that are no longer required or have become inappropriate; it is important for survival, such as stopping yourself from crossing the street when a car comes around the corner without noticing you. Deficits of response inhibition have been linked to several disorders like attention-deficit/hyperactivity, obsessive-compulsive behavior and substance abuse. Probing response inhibition has been used widely to study executive control and flexibility in behavior; for reviews, see (6–9) and a recent theme issue of *Philosophical Transactions of the Royal Society B* (2017) on ‘Movement suppression: brain mechanisms for stopping and stillness’ (10)

In the laboratory, a very useful tool for the study of inhibition is the *stop-signal paradigm* where participants perform a response time task (*go task*), such as moving their gaze to the location of a pre-defined target. Occasionally, the go stimulus is followed by a stop signal after a variable time delay, indicating subjects to withhold the response (*stop task*). Performance in the stop-signal paradigm has been modeled as a race between a “go process”, triggered by the presentation of the go stimulus, and a “stop process” triggered by the presentation of the stop signal (6, 8). When the stop process finishes before the go process, the response is inhibited; otherwise, it is executed. The main interest of the modeler is in estimating the unobservable stop-signal reaction time (SSRT), that is, the latency of the stopping process as a characterization of the response inhibition mechanism. In the *independent race model* (IND model, for short) the stop-signal task is represented as a race between *stochastically independent* go and stop processes (11). Under certain simplifying assumptions, mean SSRT can be estimated efficiently without making any specific assumptions about the distribution of the processing times (6). In several hundreds of studies, the IND model has been applied in virtually every stop-signal experiment providing important measures of cognitive control like SSRT. Although the model makes no commitment to the underlying computational or neural processes that generate the go and stop processing times, the IND model is considered as defining constraints that any model of response inhibition must follow (12).

However, neurophysiological studies in the frontal eye fields (FEF) and superior colliculus (SC) of macaque monkeys performing a countermanding task with saccadic eye movements have shown that the neural correlates of go and stop processes produce eye movement behavior through a network of *interacting* gaze-shifting and gaze-holding neurons (13–16). This discrepancy between the behavioral and neural data is widely perceived as a paradox (9, 12, 17, 18): how can interacting circuits of mutually inhibitory neurons instantiate stop and go processes with stochastically independent finishing times?

Here we propose a variant of the race model which, instead of independence, assumes *perfect negative stochastic dependence* between go and stop processes (PND model, for short). It resolves the apparent paradox and nonetheless retains the distribution-free property of the independent race model. Moreover, the PND model’s predictions, considered at the level of mean SSRT, are shown to be necessarily identical to those of the independent model. Notably, we argue that it is very difficult to empirically distinguish between the two race model versions without introducing further, parametric assumptions. Thus, there is no reason to uphold the stochastic independence assumption of the race model whenever a distribution-free version is to be considered.

## Neurally inspired modeling

Investigating the neural underpinnings of response inhibition in saccades, Hanes and Schall (13) first showed that macaque monkey behavior in saccade countermanding corresponded in detail to that of human performance in manual stop-signal tasks. Then, recording from the frontal eye fields they isolated neurons involved in gaze-shifting and gaze-holding that represent a larger circuit of such neurons that extends from cortex through basal ganglia and superior colliculus to brainstem (14). Importantly, that result was based on their postulate that for neurons to participate in controlling movement initiation two criteria must be met: first, neurons must discharge differently when movements are initiated or withheld; if neurons still discharge when movements are canceled, their activity was not affected by the stop process. Second, the differential modulation on canceled trials must occur before SSRT; otherwise, the neural modulation happens after the movement has already been canceled (18, p.1012). From these findings, Boucher and colleagues (17) developed an *interactive race model* linking the interacting circuits of mutually inhibitory gaze-holding and gaze-shifting neurons with stochastic accumulation in the go and stop processes of the race model (see also 19). A response is observed only if the go process reaches a certain threshold of activation. Stopping occurs if the stop process interferes with the go process by inhibiting activation in the go accumulator to prevent it from reaching the threshold. Alternative models assume that response inhibition results from blocking the input to the go unit (“blocked-input models”) (20) or postulate a spiking neural network of hundreds of units representing populations of movement neurons, fixation neurons and inhibitory interneurons, and a control unit that turns the fixation neurons on and off (21).

These neural models are computationally explicit: they are based on systems of stochastic differential equations; estimating parameters to best fit the behavioral data, they are able to predict the distribution of cancel times, i.e., the times at which neural activity modulates on trials on which subjects stop successfully, relative to SSRT (12). The models fit the behavioral data just as well as the independent race model. Note that this is just another instantiation of the paradox. An attempt to resolve the contradiction between stochastic independence for behavioral data and interdependence at the neural level has been to postulate that the stop process is independent of the go process for much of its duration, followed by a late and potent interaction between stopping and going that reverses the trajectory of go activation (12, 17). Strictly speaking, this would require introducing two subsequent processing stages in the IND model, independence followed by strong interaction. No formal modeling of this extension has been undertaken, however, to the best of our knowledge.

## Background: General race model

Next, we present a formal framework for the general race model making no assumption at all about the dependency between stop and go processes. One should distinguish between two different experimental conditions termed context *GO* where only a go signal is presented, and context *STOP* where a stop signal is presented in addition. A race between processes triggered by the go and stop signal is represented by two random variables: *T_go_* and *T_stop_* (referred to as SSRT above) denote the random processing times for the go and stop signal, respectively, in context *STOP* with a bivariate distribution function denoted *H*,

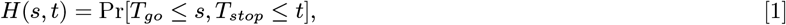

defined for all real numbers *s* and *t*, with *s*,*t* ≥ 0. The marginal distributions of *H*(*s*,*t*) are

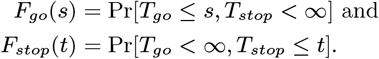

In context *STOP* the go signal triggers a realization of random variable *T_go_* and the stop signal triggers a realization of random variable *T_stop_*. In context *GO*, however, only processing of the go signal occurs. In general, the latter one may be different from the marginal distribution *F_go_*(*s*) in context *STOP*. However, the *general race model* rules this out by adding the important assumption of “context invariance”, also know as “context independence” (e.g. 9):

#### Significance Statement

The ability to suppress a response that is no longer required or has become inappropriate is an important act of cognitive control. Deficits in inhibiting a response to a stop signal have been linked to disorders like attention-deficit/hyperactivity. A widely accepted model for the stop signal paradigm assumes that stop signal processing occurs independently from go signal processing and permits to estimate the covert latency of the stop process. However, neurophysiological data suggest interactive neural networks of going and stopping raising a contradiction between the behavioral and neural findings. Introducing a model with strong negative stochastic dependency between go and stop processing resolves the paradox while retaining the computational simplicity of the original model.

### Context invariance (CI)

In context *GO*, the distribution of go signal processing time is assumed to be

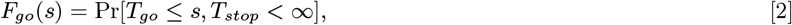

i.e., it is identical to the marginal distribution *F_go_*(*s*) in context *STOP*.

From these assumptions, the probability of observing a response to the go signal, given a stop signal is presented with delay *t_d_* [ms] (t_d_ ≥ 0) after the go signal, is defined by

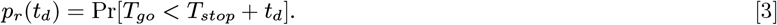

According to the model, the probability of observing a response to the go signal no later than time *t*, given the stop signal was presented with delay *t_d_*, is given by (conditional) distribution function

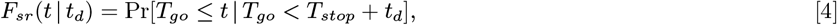

known as *signal-response RT* distribution.

The main interest in race modeling is to obtain information about the distribution of the unobservable stop signal processing time *T_stop_*, or about some of its parameters, given sample estimates of *F_go_*(*t*), *F_sr_*(*t*|*t_d_*), and *p_r_*(*t_d_*). As observed in (11), letting stop signal delay *t_d_* vary, the *inhibition function p_r_* (*t_d_*) can be formally considered as the distribution function of a random variable, *T_d_* ≡ *T_go_ − T_stop_*, say. Then,

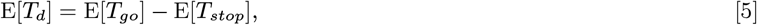

with E[] denoting expected value (mean) of a random variable. Solving for the estimate of E[*T_stop_*] immediately yields an estimate of the mean of the unobservable distribution of *T_stop_*. It is well known that the reliability of this estimation method, known as the *mean method*, depends on how precise the inhibition function *p_r_* (*t_d_*) and the mean of *T_go_* are estimated (6, 9). Obtaining estimates of higher moments, or the entire distribution of *T_stop_*, requires further non-parametric assumptions about the bivariate distribution *H*(*s*,*t*).

## Independent vs. negatively dependent race model

The most common version of the race model, as introduced by Logan & Cowan (11), postulates stochastic independence between *T_go_* and *T_stop_*:

### Stochastic independence

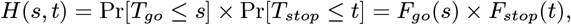

for all *s*, *t* (*s*,*t* ≥ 0).

In addition to estimating the mean via Equation [5], an estimate of the variance of stop signal processing time is obtainable in the IND model from

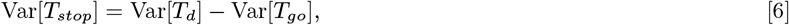

due to assuming stochastic independence. Finally, it can be shown that the unobservable stop signal distribution function *F_stop_*(*t*) is expressible as a function of the observables *p_r_*(*t_d_*), *ƒ*_*go*_(*t*) and *F_sr_*(*t*|*t_d_*) (see Methods section).

In order to resolve the paradox described above, a race model with negative dependency between go and stop signal processing times is proposed next. We define a bivariate distribution function for *T_go_* and *T_stop_* exhibiting *perfect negative dependence* (see, e.g., 22) as follows :

### Perfect negative stochastic dependence

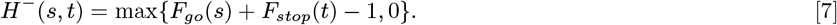

for all *s*,*t* (*s*,*t* ≥ 0). The marginal distributions of *H^−^*(*s*,*t*) are the same as before, that is, *F_go_*(*s*) and *F_stop_*(*t*). Note that this perfect negative stochastic dependence (PND) model is parameter-free just like the IND race model, that is, we do not assume some specific parametric distribution. It can be shown (see Methods) that then

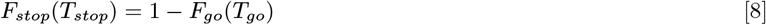

holds “almost surely”, that is, with probability 1. Thus, for any *F_go_* percentile we immediately obtain the corresponding *F_stop_* percentile as complementary probability and vice versa, expressing perfect negative dependence between *T_go_* and *T_stop_*. The relation in Equation [8] is also interpretable as “*T_stop_* is (almost surely) a decreasing function of *T_go_*”.

The PND model arguably constitutes the most direct implementation of the notion of “mutual inhibition” observed in neural data: any increase of inhibitory activity (speed-up of *T_stop_*) elicits a corresponding decrease in “go” activity (slow-down of *T_go_*) and vice versa.

### Predictions from the PND race model

We list some predictions from the PND race model (for further details, see Methods section):

#### Probability of stopping

Under perfect negative dependence between *T_go_* and *T_stop_*, the probability of stopping *p_r_*(*t_d_*) is an increasing function of *t_d_*, as it should.

#### Signal-response RT distribution

Signal-response RT distribution

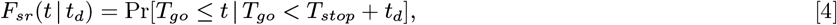

approaches the go signal distribution *F_go_*(*t*) with increasing stop signal delay *t_d_*. Moreover, for varying values of *t_d_*, *F_sr_*(*t*|*t_d_*) exhibits the “fan effect” typically found in empirical data and also predicted by the IND race model (6, 11).

To illustrate, Figure 1 presents the go signal distribution and signal-response RT distributions with different delay values *t_d_* for both the IND (dashed curves) and the PND race model (solid lines) assuming exponential distributions for *T_go_* and *T_stop_*. While the exponential distribution lacks empirical support, it was chosen here just to illustrate quantitative differences between the model versions.

**Figure caption for Figure 1.**
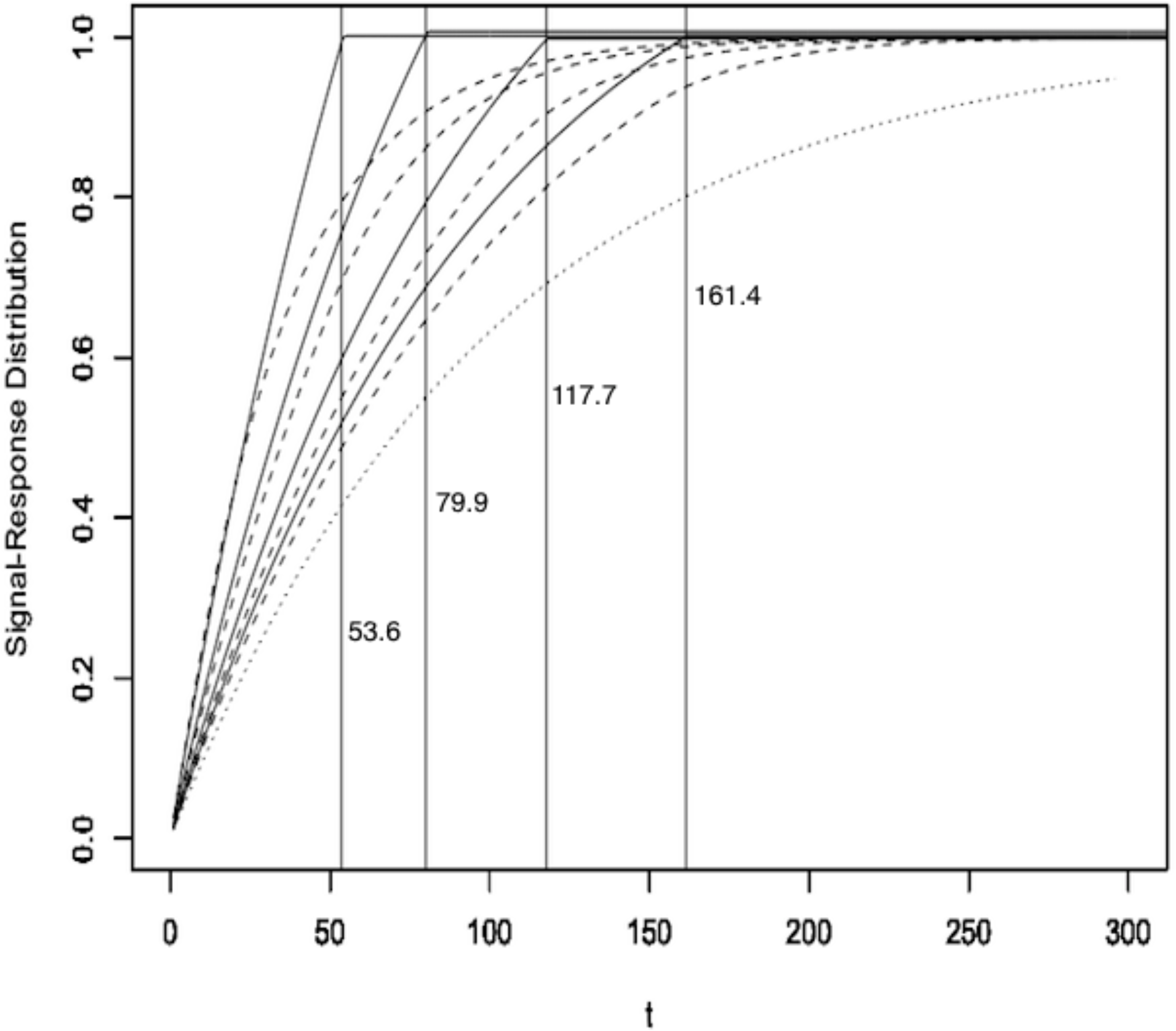
The “fan” effect: all signal-response RT distributions *F_sr_*(*t*|*t_d_*) are strictly ordered by delay (*t_d_*) from left to right, for *t_d_* = 10, 50,100,150 [ms], converging toward no-stop signal distribution *F_go_*(*t*) (rightmost curve) for *t_d_* → ∞, for exponential distributions with rate parameters *λ_go_* = .01 and *λ_stop_* = .02; PND model curves were obtained by simulation (see Methods); dashed: IND model; solid: PND model.

An important feature of the PND model is suggested by Figure 1: each signal-response distribution *F*_*sr*_ (*t*|*t_d_*) crosses 1 at a certain finite point *t*. It is shown below that this point is predictable from the observables *p_r_*(*t_d_*) and *F_go_*(*t*). While this “crossing” property is not shared by the IND model (because the “coupling” of *T_go_* and *T_stop_* as expressed in Equation (8) is absent), the strength of this test will of course depend on the precision of estimates of *F_sr_*(*t*|*t_d_*).

#### Estimating moments of the *T_stop_* distribution

Given that the marginal distributions *F_go_* and *F_stop_* are the same under both models, any estimates for E[*T_stop_*] based on the mean method (Eq. [5]) are necessarily identical for the IND and PND model.

However, for the variance we obtain

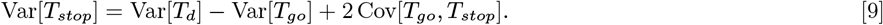

As the covariance in the above equation is unknown, an estimate for the stop signal variance is no longer available under the PND model. Nevertheless, given that Cov[*T_go_*,*T_stop_*] is the most extreme negative covariance under any bivariate distribution for *T_go_* and *T_stop_* (23), the PND model stop signal variance can never be larger than the one under independence.

## Discussion

### Testing IND vs. PND race models

Because stop signal processing times are not observable, empirical testing of non-parametric stop signal race models is severely limited in general. In particular, both the IND and PND version of the race model can only be tested in conjunction with context invariance (CI). Presuming CI is valid, the following predictions can be tested: (1) mean signal-response RT should be faster than mean go signal RT; (2) mean signal-response RT should increase with stop signal delay *t_d_*; and (3) both these tests are implied by the “fan” structure of the distribution functions permitting an additional qualitative test of IND and PND race models. Because data not consistent with this “fan” structure would be evidence against both the IND and PND model, obviously, none of these tests would allow us to distinguish between IND and PND.

By construction, the PND model has the same distribution for stop signal processing as the IND model; therefore, estimates for mean SSRT, i.e., the expected value of stop signal processing, will be identical for either model as long as estimates are based on the mean method. In a sense, this is good news because it implies that adopting perfect negative dependency between go and stop signal processing does not invalidate previous SSRT estimates of all empirical studies employing the mean method. Note that this holds as well for estimating SSRT via the *integration method*, another way to estimate SSRT. However, because the integration method presumes stop signal processing time to be constant, the distinction between IND and PND model becomes meaningless.

This leaves us with the shape of the family of signal-response distributions, that is, *F_sr_*(*t*|*t_d_*) for varying values of *t_d_*, as the only potential means of distinguishing between the two models. Whether or not this “crossing” test that -as outlined above- consists of checking the way in which the signal-response distributions approach their upper bound of 1, proves to be as visible in real data compared to the exponential toy example (Figure 1) will depend on the specific, but unobservable, stop signal distribution.

### Conclusion

Several authors have stressed that the level of description provided by the race model is quite different from that of neural models (17, 20): the independent race model is mute about the possible underlying mechanisms of response inhibition, and its primary purpose is to provide a measure of stop signal processing time, which the model allows without having to assume a specific distribution or estimate parameters. Nevertheless, it has been claimed that the independent race model “captures the essence of computation and…that it formulates the constraints that any model of response inhibition must follow” (cf. 20, p.3). The race model with perfect negative dependence suggested here operates at the same level of generality and yields the same measure of (mean) stop signal processing time but, importantly, resolves the paradox the independent model is facing due to the neurophysiological findings. We conclude that the race model with perfect negative stochastic dependence is a natural way to unify the observation of interacting circuits of mutually inhibitory gaze-holding and gaze-shifting neurons with data on the behavioral level.

Beyond the simple stop signal task, several other forms of acts of control have been studied that pose an even greater challenge for efforts to identify the underlying cognitive processes, such as switching tasks or shifting of attention (5). One promising area is *selective stopping* where response inhibition is required for some stimuli (e.g. red light) but not others (green light) (e.g., 24). Representing cognitive processes for such more complex tasks may call for types of graded, instead of perfect, dependency. Appropriate race models can easily be introduced by appealing to more general forms of stochastic dependency via copulas (22).

## Methods

This section presents some computational details for both the IND and PND model, in general and for the special case of exponential distributions (for producing Figure 1).

We assume that distribution functions *F_go_* and *F_stop_* possess densities, *ƒ_go_* and *ƒ_stop_*, and are increasing, where “increasing” always refers to “strictly increasing” here. Note that although it is usually taken for granted in the race model that densities exist, strictly speaking, this is not required neither by the independent nor the dependent race model.

### Independent race model

Under stochastic independence of *T_go_* and *T_stop_*,

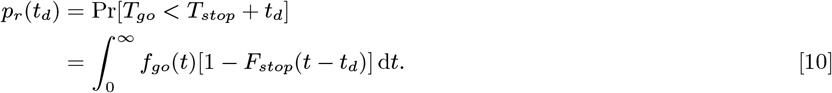

Moreover,

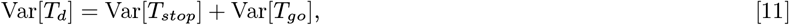

implying Equation [6].
Writing *ƒ_sr_*(*t*|*t_d_*) for the density function of signal-response time distribution *F_sr_*(*t*|*t_d_*), it has been shown in (25) that

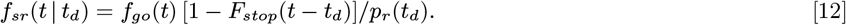

From that, an explicit expression of the distribution of the unobservable stop signal processing time *T_stop_* is given by:

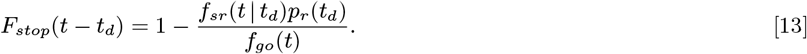

Unfortunately, as investigated in (26, 27), gaining reliable estimates for the stop signal distribution using Equation [13] requires unrealistically large numbers of observations.

### Perfect negative dependent race model

Defining the bivariate distribution by

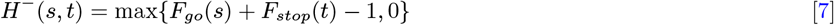

for all *s*,*t* (*s*,*t* ≥ 0), the marginal distributions of *H*^−^(*s*,*t*) are again *F_go_*(*s*) and *F_stop_*(*t*). A classic result from the *theory of copulas* (see SI) asserts that

i. *H*^−^ is the lower bound of all bivariate distributions *H* of (*T_go_*,*T_stop_*), i.e., *H*^−^(*s*,*t*) ≤ *H*(*s*,*t*) for all (*s*,*t*);
ii. Pr[*F^go^*(*T_go_*) + *F^stop^*(*T_stop_*) = 1] = 1.

From (i) the covariance between the random variables can be shown to be the smallest possible across all bivariate distributions *H*(*s*,*t*) (23). Statement (ii) is equivalent to

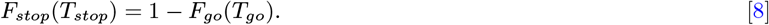

holding almost surely.

### Probability of stopping

Under perfect negative dependence between *T_go_* and *T_stop_*,

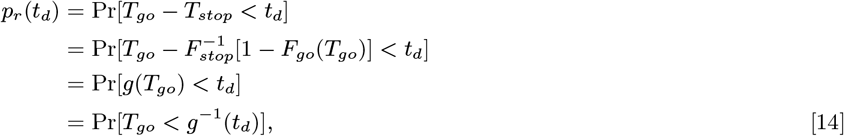

where function *g*, defined as 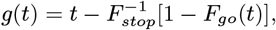, is increasing and so its inverse *g*^−1^ is increasing as well. Thus, *p_r_*(*t_d_*) is increasing in *t_d_*.

### Signal-response RT distribution

For stop signal delay *t_d_*,

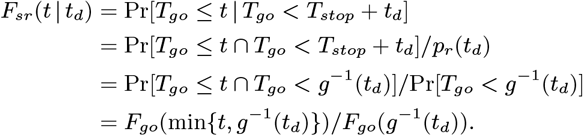

To see that *F_sr_*(*t*|*t_d_*) obeys the “fan effect” consider two different delays, *t_d_* and 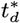, say, with *t_d_* < 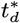; for a fixed value of *t*, we first assume *t* < *g*^−1^(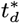) Then

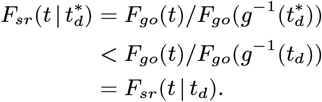

The cases *g*^−1^(*t_d_*) < *t* < *g*^−1^(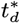) and *g*^−1^(*t_d_*) < *g*^−1^(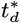) < *t* can be shown similarly.

### Expected stop signal processing time

A formal expression for mean (expected) stop signal processing time is obtained as follows. We have

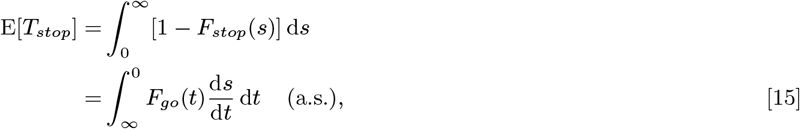

using Equation [8] and the fact that 1 – *F_stop_* (*s*) and *F_go_*(*t*) have opposite limit values. With

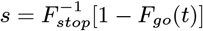

we have

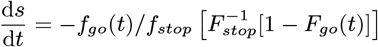

Inserting into [15] yields

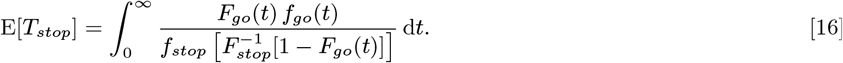

### IND model: exponential case

Define independent, exponential distributions for *T_go_* and *T_stop_* with parameters *λ_go_* > 0 and λ_*stop*_ >0 for context *STOP* by

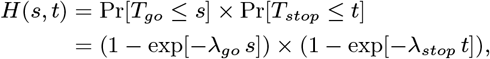

for all *s*,*t* ≥ 0. Then

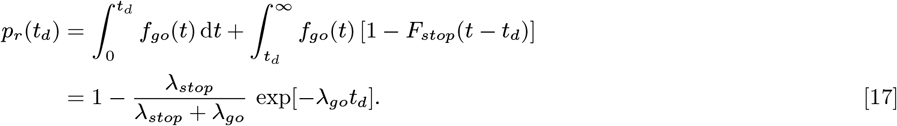

For *t* > *t_d_*, the density of the signal-response distribution is given by,

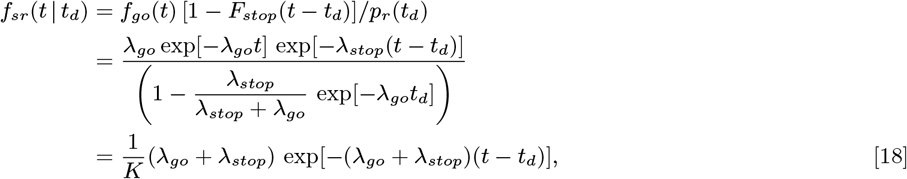

with *K* = exp[λ_*go*_ *t_d_*](1 + *λ_stop_/λ_go_) − λ_stop_/λ_go_*. For *t_d_* = 0, we have *K* = 1 and the signal-respond density is identical to an exponential density for an independent race between *T_stop_* and *T_go_*, with parameter *λ_go_ + λ_stop_* and *p_r_*(*t_d_*) = λ_*go*_/(λ_*go*_ + λ_*stop*_). For *t* ≤ *t_d_*, the density simplifies to

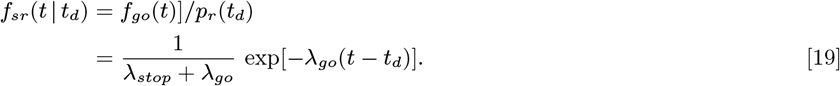

Computation of the expected value of signal-response RTs yields:

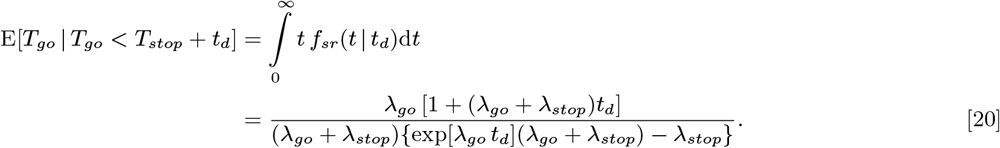

In particular, for *t_d_* = 0, we obtain E[*T_go_*|*T_go_* < *T_stop_* + *t_d_*] = 1/(λ_*go*_ + λ_*stop*_), consistent with the density we mentioned above for this value of the stop signal delay.

### PND model: exponential example

Inserting exponential margins into the bivariate distribution,

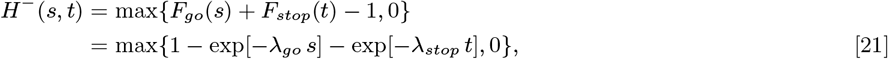

for all *s*,*t* ≥ 0. From [14],

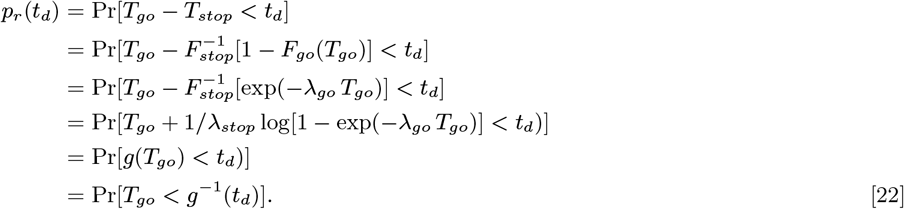

Note that function

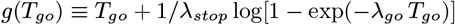

cannot be solved explicitly for *T*_go_. Therefore, in order to compute *p_r_* (*t_d_*) and plot signal-response time distributions

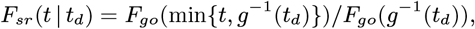

we sampled (*n* = 100, 000) from the bivariate distribution function *H*^−^(*s*,*t*) using function simCop (based on the conditional simulation method, see (22)) from the copBasic package of the open source software R (http://www.r-project.org).

Table 1 lists the crossing points for the signal-response time distributions obtained from the simulation. They correspond to the vertical lines in Figure 1.

**Table 1.** Predictions for crossing points *F_sr_*(*t*|*t_d_*) = 1.0 in PND model. Compare with Figure 1

| $t_d$ [ms] | 10 | 50 | 100 | 150 |
| --- | --- | --- | --- | --- |
| $g^{-1}(t_d)$ | 53.6 | 79.9 | 117.7 | 161.4 |

## Supporting Information (SI)

### Fréchet-Hoeffding bounds and perfect dependency

The dependence between two random variables of a random vector (*X*, *Y*) L is completely described by its probability distribution. Let *G*(*x*,*y*) be the bivariate distribution function of some random vector ! (*X*,*Y*):

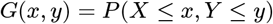

with marginal distributions *F_X_* and *F_Y_*. Then, it always holds that

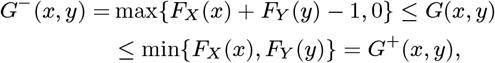

for all *x*,*y* for which *G* is defined (the support of *G*). Both *G*^−^ (*x*,*y*) and *G*+(*x*,*y*) are known as *Fréchet-Hoeffding bounds* and are themselves distribution functions for (*X*,*Y*): *G*^−^ corresponds to “perfect” negative dependence between X and *Y*, while *G+* corresponds to “perfect” positive dependence (see (22) for proofs of this and other claims in this section). “Perfect” dependency can be formulated in various ways. For simplicity, we assume *X* and *Y* have continuous distribution functions, the only case we will need below. Here are three equivalent characterizations of *perfect negative dependence* (analogous results exist for perfect positive dependence, but they are not of concern here):

i. Either Pr(*X* > *x*,*Y* > *y*) = 0 for all *x*,*y* or Pr(*X*≤ *x*,*Y* ≤ *y*) = 0 for all *x*, *y*;
ii. Pr[*F_x_* (*X*) + *F_Y_*(*Y*) = 1] =1;
iii. *X* is almost surely a decreasing function of *Y*.

Consider the equation inside the “Pr” expression in (ii): Solving for *X* by taking the inverse function of *F_x_*(*X*) we get

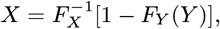

from which (iii) follows. Random variables with perfect negative dependence are also known as *antithetic variates* in simulation studies.

